# *propeller*: testing for differences in cell type proportions in single cell data

**DOI:** 10.1101/2021.11.28.470236

**Authors:** Belinda Phipson, Choon Boon Sim, Enzo R. Porrello, Alex W Hewitt, Joseph Powell, Alicia Oshlack

## Abstract

**Motivation:** Single cell RNA Sequencing (scRNA-seq) has rapidly gained popularity over the last few years for profiling the transcriptomes of thousands to millions of single cells. This technology is now being used to analyse experiments with complex designs including biological replication. One question that can be asked from single cell experiments, which has been difficult to directly address with bulk RNA-seq data, is whether the cell type proportions are different between two or more experimental conditions. As well as gene expression changes, the relative depletion or enrichment of a particular cell type can be the functional consequence of disease or treatment. However, cell type proportions estimates from scRNA-seq data are variable and statistical methods that can correctly account for different sources of variability are needed to confidently identify statistically significant shifts in cell type composition between experimental conditions.

**Results:** We have developed *propeller*, a robust and flexible method that leverages biological replication to find statistically significant differences in cell type proportions between groups. Using simulated cell type proportions data we show that *propeller* performs well under a variety of scenarios. We applied *propeller* to test for significant changes in proportions of cell types related to human heart development, ageing and COVID-19 disease severity.

**Availability and implementation:** The *propeller* method is publicly available in the open source `speckle` R package (https://github.com/phipsonlab/speckle). All the analysis code for the paper is available at https://github.com/phipsonlab/propeller-paper-analysis/, and the associated analysis website is available at https://phipsonlab.github.io/propeller-paper-analysis/.

**Contact:** Alicia Oshlack

Belinda Phipson

**Supplementary information:** Yes.

## 1. Introduction

Single cell RNA-sequencing (scRNA-seq) technology has led to breakthroughs in the discovery of novel cell types and enhanced our understanding of the development of complex tissues. As the technology has matured it has become relatively straightforward to profile the transcriptomes of hundreds of thousands of cells, resulting in valuable insight into the composition of tissues.

While many of the first published single cell papers focused on defining the resident cell types in complex tissues^1–4^, the field is now using this technology for complex experimental comparisons with biological replication^5–8^. Experiments with different conditions and multiple biological samples can be costly, however substantial savings can be made by pooling cells from multiple samples. If samples are genetically diverse, they can be demultiplexed using genetic information^9,10^. An alternative approach is to use molecular cell multiplexing protocols, such as the commercially available CellPlex from 10x Genomics. Collectively, cell multiplexing makes designing larger scRNA-seq experiments more feasible.

The first step in analysis for an scRNA-seq experiment with multiple experimental conditions and biological replicates is to identify the cell types present in each sample. However, downstream analysis requires sophisticated tools to address specific hypotheses about how a perturbation affects the biological system. Two analysis tasks are commonly performed following cell type identification in order to understand the effect of the condition. One task is to find genes that are differentially expressed between groups of samples, for every cell type observed in the experiment, similar to the analysis of bulk RNA-seq experiments^11^. However, a benefit of scRNA-seq data is that we have additional information on the composition of the samples. The relative change in abundance of a cell type can be a consequence of normal development, disease or treatment. Due to technical as well as biological sources, the cell type proportions estimates from single cell data can be quite variable. The focus of this work is to find statistically significant differences in cell type proportions between groups of samples that appropriately takes into account sample-to-sample variability.

Here we present *propeller*, a robust and flexible linear modeling based solution to test for differences in cell type proportions between experimental conditions. The *propeller* method leverages biological replication to obtain measures of variability of cell type proportion estimates, and uses empirical Bayes to stabilise variance estimates by borrowing information across cell types. It is a flexible approach that can be applied to complex experimental designs with multiple factors. Using simulated data, we compared the performance of commonly used statistical models for testing for differences in cell type proportions and show that *propeller* performs well across a variety of experimental scenarios. We applied *propeller* to three different single cell datasets on ageing, human heart development and COVID-19 disease severity and find additional cell types changing in abundance that were not reported in the original analysis. Our *propeller* method is publicly available in the `speckle` R package (https://github.com/phipsonlab/speckle).

## 2. The *propeller* method

*Propeller* is a function in the `speckle` R package that uses cell level annotation information to calculate sample level cell type proportions, followed by data transformation and statistical testing for each cell type. *Propeller* leverages biological replication to estimate the high sample-to-sample variability in cell type counts often observed in real single cell data (Figure 1a, PBMC scRNA-seq data from 12 healthy human donors). The variability in cell type proportions estimates between samples can be large both due to technical sources such as variation in dissociation protocols, and due to valid biological factors that contribute to variability. For example, blood cell type composition is known to change with age^12^. Taking into account sample-to-sample variability when analysing differences in cell type proportions is critical as observed cell type variances are far greater than variances estimated under a binomial or Poisson distribution, which can only account for sampling variation (Figure 1b, PBMC scRNA-seq dataset from 12 healthy human donors).

**Figure 1.**
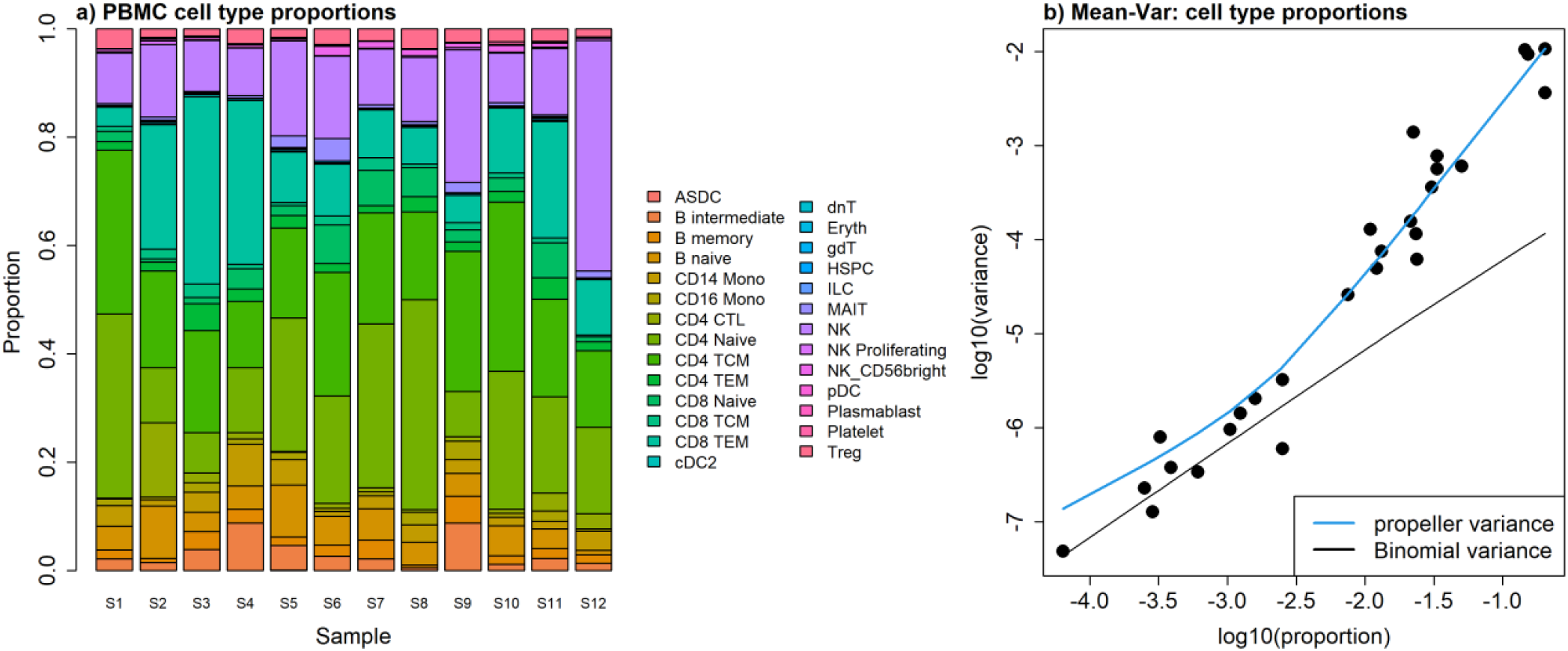
Exploring heterogeneity in cell type proportions estimated from single cell RNA-seq data. **a**. Barplot showing high levels of variability of cell type proportion estimates between 12 healthy PBMC scRNA-seq samples. **b**. Mean-variance relationship for 27 cell types in 12 healthy PBMC scRNA-seq samples showing that cell type proportions are over-dispersed compared to the variance estimated under a Binomial distribution. The plot is produced using the *plotCellTypePropsMeanVar* function in the `speckle` package.

The first step of *propeller* is to calculate the cell type proportions for each sample. *Propeller* can directly derive the counts and calculate the proportions from a Seurat or SingleCellExperiment object. This results in a matrix of proportions where the rows are the cell types and the columns are the samples. The binomial distribution has the statistical property that proportions close to 0 and 1 have small variance, and values close to 0.5 have large variance i.e. the variances are heteroskedastic. To overcome this, we have implemented two transformations in *propeller*: (1) arcsin square root transformation, and (2) logit transformation. The arcsin square root transformation has the benefit that it will always produce a real value. If the logit transformation is selected an offset of 0.5 is added to the raw cell type counts matrix prior to transformation to avoid taking the log of zeroes.

Next we test whether the transformed proportions for every cell type are significantly different between two or more experimental conditions using a linear modelling framework. If there are exactly two groups, we perform moderated t-tests; if there are more than two groups, we perform moderated ANOVA tests^13^. These tests are moderated using an empirical Bayes framework, allowing information to be borrowed across cell types to stabilise the cell type specific variance estimates. This is particularly effective when the number of biological replicates is small and the number of cell types is at least three^14^, a common situation in scRNA-seq experiments. The final step in *propeller* is to calculate false discovery rates^15^ to account for testing across multiple cell types. The output of *propeller* consists of condition specific proportions estimates, *p*-values and false discovery rates for every cell type observed in the experiment.

The minimal annotation information that *propeller* requires for each cell is cluster/cell type, sample and group/condition, which can be manually entered or automatically extracted from Seurat and SingleCellExperiment class objects. More complex experimental designs can be accommodated using the *propeller*.*ttest* and *propeller*.*anova* functions, which have the flexibility to model additional covariates of interest such as sex or age.

## 3. Performance using simulated datasets

### 3.1 Type I error control under null simulation scenario

Although it is clear from the PBMC scRNA-seq data that cell type proportions estimates are overdispersed (Figure 1b), we wanted to more thoroughly evaluate the performance of *propeller* as well as other statistical methods that have commonly been used for testing differences in proportions in other fields. Using simulated cell type proportions, we compared the performance of nine different statistical models.

1. Chi-square test for differences in proportions
2. Logistic binomial regression (special case of beta-binomial with dispersion=0)
3. Poisson regression (special case of negative binomial with dispersion=0)
4. *propeller* with arcsin square root transformation of proportions, denoted propeller(asin)
5. *propeller* with logit transformation of proportions, denoted propeller(logit)
6. Beta-binomial regression on cell type counts
7. Negative binomial regression on cell type counts
8. Quasi-likelihood negative binomial regression on cell type counts
9. Centred log-ratio transformation (CLR) followed by linear regression, denoted CODA

The first three methods do not take into account sample to sample variability, while the remaining 6 methods (*4-9*) do. The quasi-likelihood approach (method 8) is described in the Bioconductor book “Orchestrating single cell analysis with Bioconductor”^16^. The two variations of *propeller* model transformed proportions, while the remaining statistical tests, with the exception of the CODA method, model the cell type counts directly. Method 9 is an example from the compositional data analysis field where the cell types are modeled relative to a reference “cell type”. Here the geometric mean of the cell types forms the baseline as is commonly done in microbiome data analysis^17^. The log-ratio of the counts to the geometric mean is calculated and a linear model fitted with group as the explanatory variable to obtain p-values.

We simulated cell type counts in a hierarchical manner under a simple null scenario where the cell type proportions do not differ between two groups in order to determine whether the nine methods control the Type I error rate. We simulated five cell types with proportions that varied from rare to abundant (true proportions *π*_*i*_ = 0.01, 0.02, 0.15, 0.34, 0.45). The sample proportions, *p*_*ij*_, for cell type *i* and sample *j*, were generated from a Beta distribution with parameters *α*_*i*_ and β_*i*_, which control how variable the proportions are. Larger values of *α*_*i*_ and β_*i*_ result in a more precise distribution centred around the true proportions, while smaller values result in a more diffuse prior (Supplementary Figure 1). We set *α*_*i*_ = 10, and calculated the corresponding *β*_*i*_ for each cell type *i* from the following relationship derived from properties of the Beta distribution:

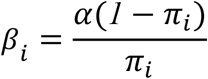

Cell type counts *x*_*ij*_ were then sampled from a binomial distribution with parameters *n*_*j*_ and *p*_*ij*_. The total number of cells, *n*_*j*_, per sample *j*, were sampled from a negative binomial distribution with mean 5000 and dispersion 20 to simulate variation in total cell numbers per sample observed in real data. We simulated 10,000 datasets, and counted the number of times each of the five cell types were statistically significant with p-value < 0.05 for the nine different statistical models. We also varied the number of samples per group to determine the effect of sample size on the Type I error rate (*n* = 3, 5, 10, 20). Figure 2a shows the cell type proportions per sample observed for an example simulated dataset under these conditions.

**Figure 2.**
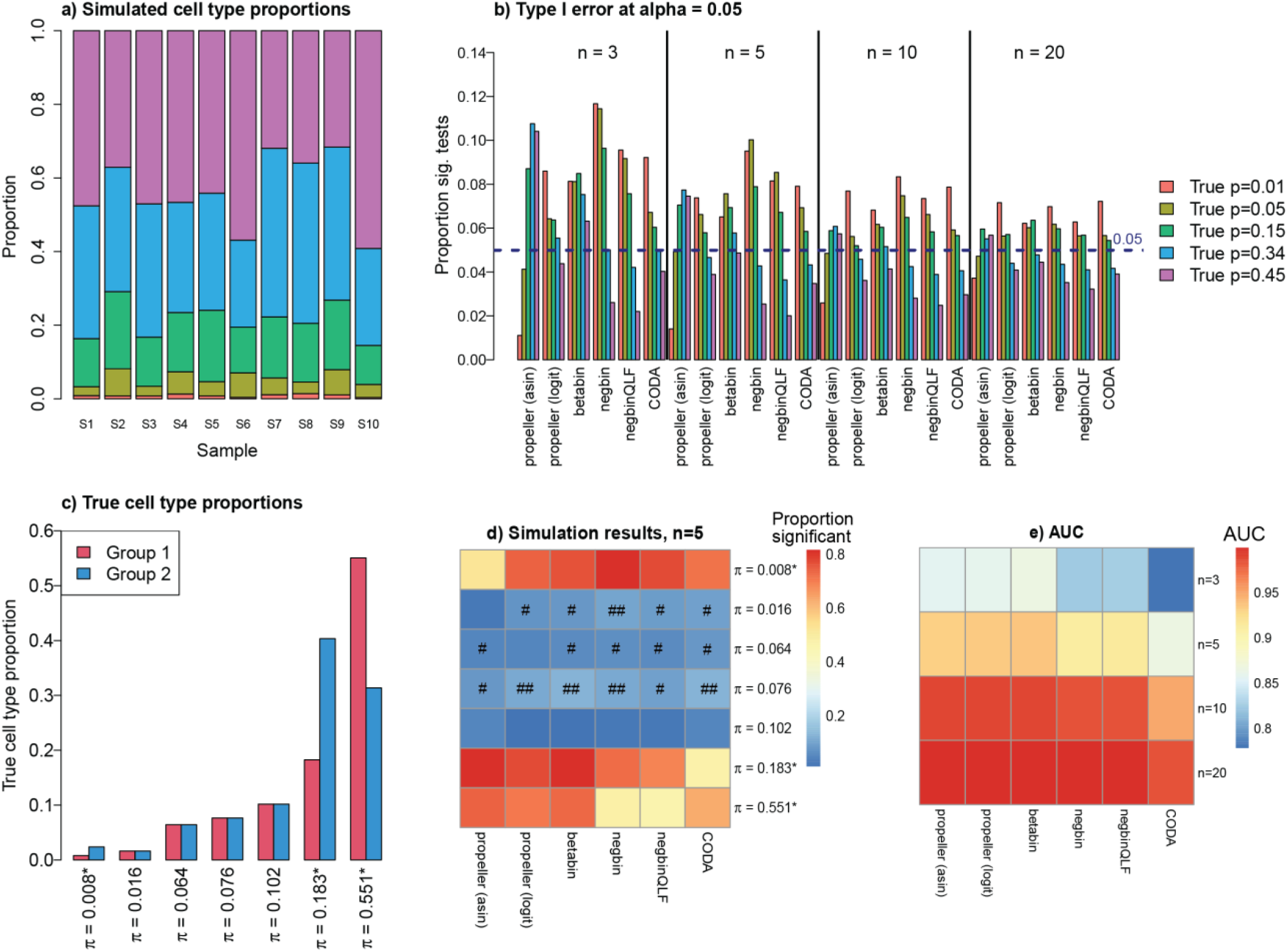
Simulation results. **a**. Cell type proportions for one simulated dataset with no abundance differences between group 1 (samples S1 - S5) and group 2 (samples S6 - S10). **b**. Type I error rate at *α*=0.05 for sample sizes n=3, 5, 10, 20 for the six methods. 1000 datasets with 5 cell types that do not change in abundance between two groups were simulated, varying the sample size. For each of the cell types, the proportion of simulated datasets with p-value < 0.05 was calculated when testing for cell type proportion differences for each of the six models. **c**. True cell type proportions for group 1 and group 2. Three cell types that range in abundance are simulated to vary by 2 - 3 fold (denoted by asterisks). The remaining four cell types do not differ. **d**. Heatmap showing the proportions of 1000 simulated datasets with p-value < 0.05 when testing for cell type proportion differences between two groups. True positives are denoted by an asterisk. Dark red indicates greater power to detect significant cell type differences (proportion significant is high). For true negatives, dark blue without the # symbol indicates good false discovery rate control with proportion significant <0.05, # indicates proportion significant between 0.05 and 0.1, and ## indicates poorest control with the proportion significant > 0.1. **e**. Heatmap showing the mean area under the receiver operating curve (AUC) for each of the six methods for all sample sizes across 1000 simulated datasets. Higher AUC (darker red) indicates a method has both good power to detect true positives as well as good false discovery rate control.

Table 1 shows the type I error rates for the nine different methods for each of the five different cell types at a nominal p-value cut-off of 0.05 when the number of samples per group is five. The most striking observation is that the statistical tests (methods *1-3*) that do not account for additional biological variability frequently find significant differences between the two groups when there are none. As expected, it is clear that methods that account for this additional variability are required and methods *1-3* are not further explored in this analysis.

**Table 1.**
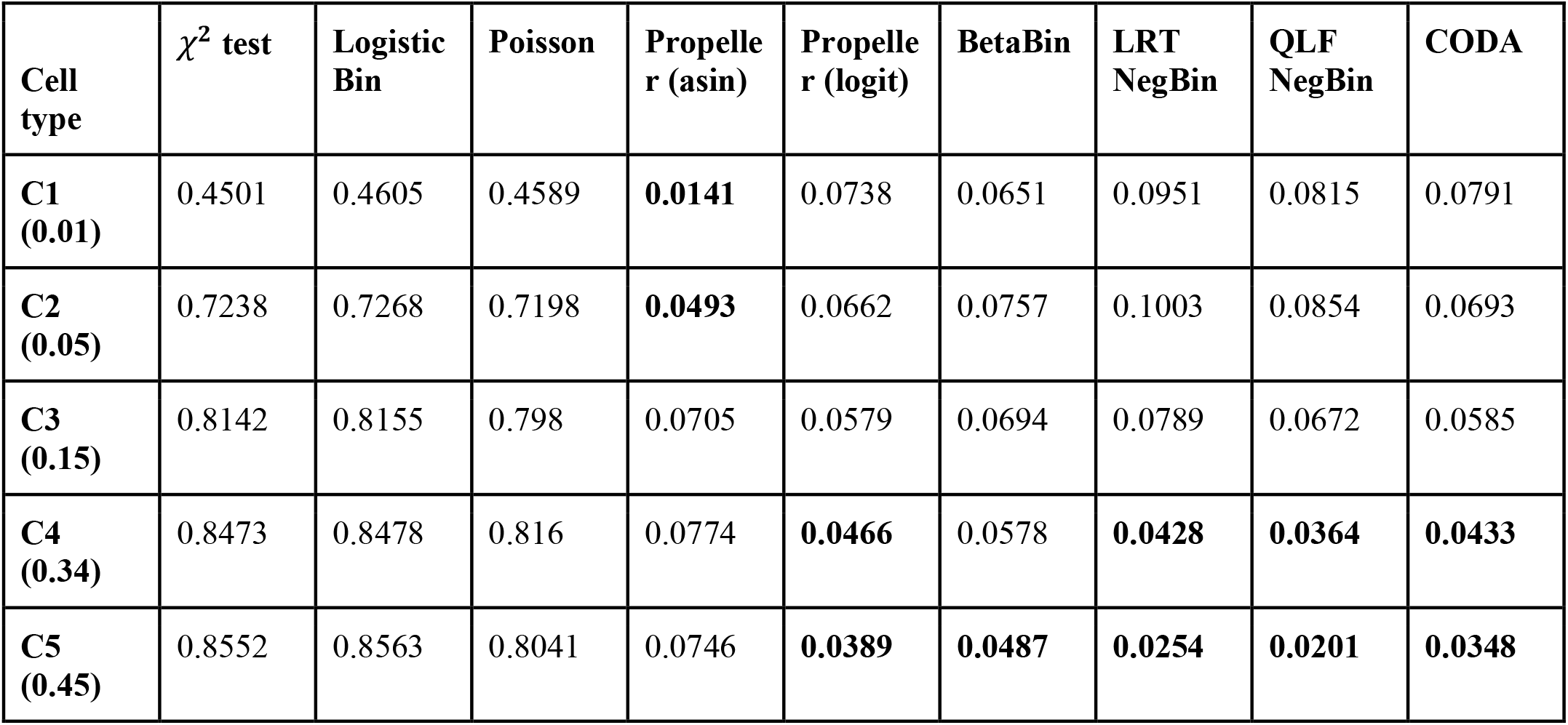
Proportion of significant tests for each cell type between two groups with no differences at a nominal p-value cut-off of 0.05. Ten thousand datasets were simulated under a beta binomial model with five samples in each group. In order to control the type 1 error rate correctly, the proportion of significant tests should be approximately 0.05 for each cell type. The number in brackets in the first column corresponds to the true cell type proportion, and the cell types are ordered from most rare to most abundant. The bold numbers indicate Type I error estimates < 0.05.

For the methods that model sample-to-sample variability none have perfect type I error rate control, although the observed rates are generally close to 0.05 (methods *4-9*). Propeller(asin) tends to be conservative for the most rare cell type, and permissive for more abundant cell types whereas the opposite tends to be true for the other tests, particularly for the negative binomial methods. These results show that the type I error rate differs between different cell types depending on how abundant the cell type is, and no method perfectly controls the type I error rate for both rare and abundant cell types.

Figure 2b summarises the Type I error rates for different sample sizes. As the number of samples in each group increases, the type I error rate for all methods is closer to 0.05 (Figure 2b). For sample sizes of 10 and 20 per group, the arcsin-square root transformation shows the best type I error rate control for almost all cell types abundances, however with smaller sample sizes (*n* = 3, 5), the logit transform appears to better control the type I error rate. Across all sample sizes there was a trend of increased Type I error rate for less abundant cell types for propeller(logit), beta-binomial, negative binomial, quasi-likelihood negative binomial and the CODA method, while propeller(asin) tends to be conservative for the most rare cell type (*π* = 0.01). It is not surprising that the beta-binomial model performs favourably as this method most closely resembles the distributional assumptions underlying the simulation.

### 3.2 Power to detect true differences in cell type proportions in simulated data

Next we expanded the simulation to include seven cell types, three of which change proportions between the two groups by between 2 and 3-fold, while the remaining four did not (Figure 2c). The parameters α_*i*_ and β_*i*_ of the beta distribution were estimated from real human heart single nuclei RNA-seq data (Figure 3a) using the *estimateBetaParamsFromCounts* function available in the `speckle` package. We simulated 1000 datasets and evaluated the performance of the models by examining the proportion of simulated datasets with p-value < 0.05 for each of the seven cell types for each of the six methods. The proportions of the three cell types that are simulated to differ between the two groups range from very rare to quite abundant (baseline proportions in group 1 = 0.008, 0.183, 0.551). We repeated these simulations for sample sizes *n =* 3, 5, 10, 20.

**Figure 3.**
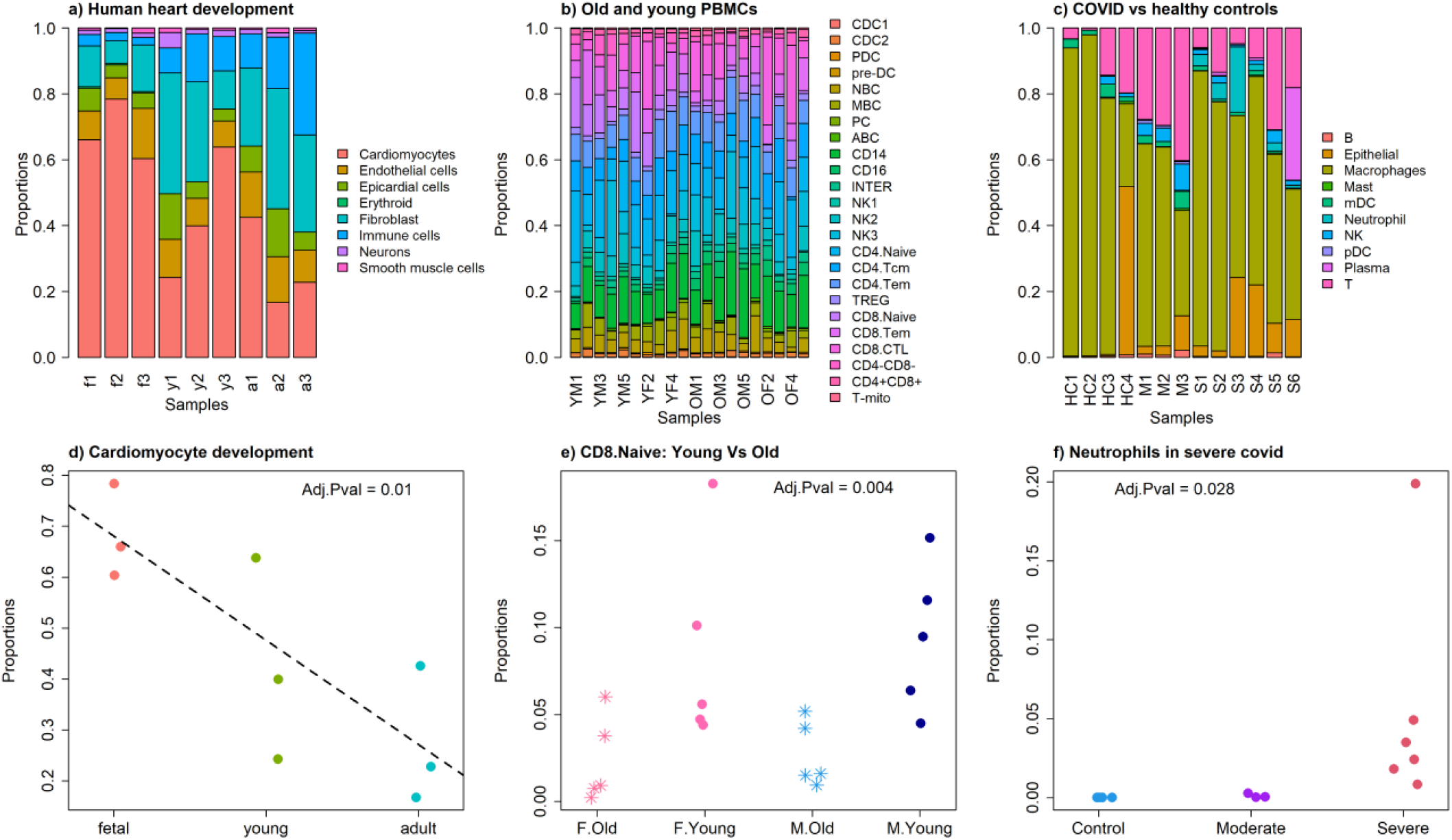
Applying propeller to three single cell RNA-seq datasets. **a**. Barplot showing cell type proportions for nine samples in a human heart development snRNA-seq dataset. f=fetal, y=young, a=adult **b**. Barplot showing cell type proportions for 20 PBMC samples that differ in terms of their age (Y/O) and sex (M/F) **c**. Barplot showing cell type proportions for 13 samples in a COVID-19 study. HC=healthy control, M=moderate COVID-19 infection, S=severe COVID-19 infection. **d**. Treating developmental stage as a continuous variable, the cardiomyocyte populations show a relative decline across development in human heart samples. **e**. There is a statistically significant difference in the proportions of CD8 naive cells between young and old PBMC samples, taking sex into account. **f**. Neutrophils are statistically significantly different between healthy control, moderate and severe COVID-19 bronchoalveolar lavage samples.

At *n=5* samples per group, propeller(asin) detected the rare cell type difference in only 52% of the simulated datasets, while the other methods detected the rare cell type difference in 71-81% of simulated datasets (Figure 2d). However, propeller(asin) detected the differences in the more abundant cell types in a larger percentage of the simulated datasets compared to the other methods (82% and 74% of simulated datasets). The negative binomial methods detected the difference in the most abundant cell type in < 50% of the simulated datasets. The CODA method had relatively poor performance for the more abundant cell types compared to the propeller methods and the beta-binomial model. The most consistent performing models across cell type abundances were propeller(logit) and the beta-binomial model. In terms of the cell types that did not change between the two groups, we noted that propeller(asin) generally had the best false discovery rate control and CODA had the worst. Heatmaps for sample sizes *n* = 3, 10 and 20 are shown in Supplementary Figure 2.

Figure 2e shows the mean area under the receiver operating curve (AUC) for the six methods at the four different sample sizes (*n* = 3, 5, 10, 20). As sample size increases, all methods show an improvement in performance. With at least 10 samples in each group, all methods except CODA have an AUC above 98%. In general, propeller(asin), propeller(logit) and the Beta binomial method have the highest AUC at each of the four sample sizes.

### 3.3 Extreme case: varying numbers of cell types

While the simulations above examine Type I error control and power to detect true positives with 5 and 7 cell types respectively, we wanted to examine the performance of the methods in the extreme case when there are only 2 cell types present in the dataset, compared to when there are 20. Here we focussed on *n* = 5 and simulating cell types with true differences between two groups.

The scenario when only two cell types are present in the data is interesting from the perspective that if one cell type changes in proportion, the other cell type will also naturally change. In this scenario, we set the Group 1 true proportions as *π*_*1i*_ = 0.4, 0.6; and Group 2 true proportions as *π*_*2i*_ = 0.2, 0.8 (Supplementary Figure 3a). Hence, cell type 1 is halved in Group 2 compared to Group 1, and cell type 2 increases by 33.3%. In this scenario, all the methods detected the changes in the two cell types in the majority of the simulated datasets (Supplementary Figure 3b).There was a slight decrease in power for the negative binomial methods for the more abundant cell type.

For the scenario with 20 cell types, we used cell type proportion estimates from the 12 healthy human PBMC scRNA-seq dataset as our true baseline proportions. We modified 8 of the 20 cell types to be different between the two groups (Supplementary Figure 4a). The cell types that differed in abundance between the two groups ranged from rare to abundant. The heatmap in Supplementary Figure 4b shows the proportion of significant tests across the 1000 simulated datasets for each cell type and each method. In this scenario with a larger number of cell types, the negative binomial methods have similar performance compared to the other methods. Cell types with larger log-fold changes are detected as statistically significant in the majority of simulated datasets by all methods.

Table 2 shows the recall, precision and F1 score for each method averaged across the 1000 simulated datasets. In this scenario, the CODA method is able to detect more true differences in cell type proportions compared to any of the other methods, at the expense of detecting the most false positives. In general, the propeller methods have high precision indicating that not many false discoveries are reported. The negative binomial methods perform well in this scenario, and the beta-binomial model has the second highest F1 score with a good balance between precision and recall. Propeller(logit) has the highest precision and CODA has the highest recall.

**Table 2.** Recall, precision and F1 score for 1000 simulated datasets with 20 cell types. 8 out of 20 cell types differ in abundance between two groups by between 1.4 - 3 fold. Recall, precision and F1 scores were calculated for each of the 1000 simulated datasets, and mean values shown. Bold values highlight best performing method for each metric.

| Method | Recall | Precision | F1 score |
| --- | --- | --- | --- |
| propeller(asin) | 0.646125 | 0.9085725 | 0.7463168 |
| propeller(logit) | 0.694250 | **0.9161798** | 0.7808882 |
| Beta-binomial | 0.725125 | 0.9091364 | 0.7984171 |
| Negative binomial | 0.710500 | 0.8972013 | 0.7839388 |
| Quasi-likelihood neg bin | 0.690125 | 0.9098287 | 0.7756611 |
| CODA | **0.794500** | 0.8640721 | **0.8197626** |

## 4. Application to real single cell datasets

One important feature of *propeller* is that complex experimental designs can be modelled by using a design matrix that takes account of multiple factors. In order to demonstrate the types of experimental designs that can be accommodated, we applied propeller(logit) to three different scRNA-seq datasets that varied in terms of the experimental design and the number of samples and cell types in each dataset:

1. 9 human heart biopsy samples across development (fetal, young, adult), with 8 broad cell types annotated^5^. We modelled development as a continuous variable and sex as a categorical variable.
2. 20 PBMC samples across young and old male and female samples with 24 cell types annotated^6^. We modelled age and sex as categorical variables.
3. 13 bronchoalveolar lavage fluid immune cell samples across three groups (healthy controls, moderate and severe COVID-19 infection) with 10 cell types annotated^18^. We modelled disease status as a categorical variable and performed an ANOVA to find cell type differences between the three groups.

Figure 3a-c shows the cell type proportion estimates for each sample for the three different datasets. The cell type proportions are highly variable between individuals across all datasets. Across healthy human heart development, we detected significant changes in the abundances of immune, erythroid, cardiomyocyte and fibroblast cells (Supplementary Figure 5). In the original analysis, propeller(logit) was applied as an ANOVA test, ignoring sex. While the conclusions are not markedly different, the order of significant cell types has changed with immune cells the most significant cell type when modelling development as a continuous variable. The immune and erythroid cell type changes across development form a type of positive control and it is encouraging that they are the most significant cell types. As noted in the initial paper^5^, immune cells increase throughout development, as would be expected as the fetus has not been exposed to many pathogens, while an adult would have a larger and more diverse repertoire of immune cells. With the erythroid cells, only fetal red blood cells are nucleated, and hence they are captured with the nuclei protocol in fetal samples and absent in young and adult samples. An interesting finding in this study was that the relative abundance of cardiomyocytes decline with age (Figure 3d), while fibroblasts increase across development (Supplementary Figure 6).

For the aging PBMC dataset, we detected statistically significant differences in CD8 naive and CD16 cells between young and old samples, while controlling for sex, at a false discovery rate threshold of 0.05. CD8 naive cells were enriched in the young samples (Figure 3e), and CD16 cells were depleted in the young samples compared to old (Supplementary Figure 6). While the CD8 naive result was reported in the initial paper, we detected a significant change in abundance of CD16 cells between young and old samples that was not reported in the original analysis^6^.

For the COVID-19 dataset, we found four cell types to have significant changes in abundance between the three groups when we used propeller(logit) (Supplementary Figure 7). We found that neutrophils were the most significantly different between healthy controls and moderate and severe bronchoalveolar lavage samples from COVID-19 patients (Figure 3f), and this was not reported as statistically significant in the original analysis^18^. Plasma, pDC and NK cells also showed significant changes in abundance. Upon closer inspection, it appeared that the significant result for Plasma was driven by one sample in the severe COVID-19 group (Supplementary Figure 7). When we re-analysed the data with propeller(asin), this cell type was no longer significant, while neutrophils, pDC and NK cells were still statistically significant. This indicates that propeller(logit) may be sensitive to outlier samples and suggests that propeller(asin) is a more robust method to use when outliers are present in the data. Compared to the results from the original analysis, we detected two additional cell types, neutrophils and NK cells, that significantly changed in abundance between healthy controls, moderate and severe COVID-19 patients.

## 5. Discussion

In this paper we present *propeller*, a new method for testing for differences in cell type proportions from single cell data. It takes account of sample-to-sample variability, which is large due to both technical and biological sources. The *propeller* function itself inter-operates with Seurat and SingleCellExperiment class objects, and the *propeller*.*ttest* and *propeller*.*anova* functions have the ability to model complex experimental designs. In order to work specifically with the features of single cell data which often have extreme cell proportions, we have implemented *propeller* with two different transformations: the arcsin square root transformation and the logit transformation. Through simulation studies, we found propeller(logit) has superior performance in terms of power to detect changes in cell type proportions, as well as good false discovery rate control. Through analysis of real datasets, we found that propeller(logit) may be sensitive to outlier samples, while propeller(asin) is not, which suggests that propeller(asin) is a good alternative in this scenario. A recent comparison of statistical methods for performing cell type composition analysis of single cell data found that propeller(asin) and Dirichlet regression had the best performance across a variety of scenarios^19^. The *propeller* methods have the ability to handle zeros and ones in the data, which are not uncommon in cell type proportion estimates from single cell data. Zero values need to be carefully dealt with when using compositional data analysis methods (CODA). For the simulation studies, we replaced zeroes with 0.5 prior to centred-log ratio transformation. Another factor to consider when using a CODA framework is the choice of reference cell type, and all results need to be interpreted relative to the reference cell type, which can make interpretation of the output more challenging.

In our simulation studies we explored the effect of the number of cell types on the performance of the methods. For datasets with fewer cell types, the negative binomial methods and the CODA method show decreased performance compared to beta-binomial and *propeller* methods. As the number of cell types increases to 20, the performance of negative binomial and CODA methods improve to be comparable to the other methods. We also explored the effect of sample size and baseline abundance of the cell type on the performance of the methods. For small sample sizes the differences between the methods is more pronounced, with propeller(logit) and beta-binomial showing the best overall performance. As the sample size increases beyond 10 samples per group, all methods show good power and false discovery rate control, with the exception of the CODA method, which has increased false discovery rates for all cell types with increasing sample size. We also found that at smaller sample sizes, propeller(asin) had less power to detect the differences in the most rare cell types, while the negative binomial methods had less power to detect differences in the most abundant cell types.

We applied *propeller* to the analysis of three different single cell datasets that differed in terms of tissue, number of cell types, sample size and experimental conditions. We found significant biological differences in abundance, including some cell types that had not been previously reported, in three different studies: across healthy human heart development, comparing blood from young and old patients, and in lung fluid from individuals with severe covid versus moderate and healthy controls. All our analysis is available via a *workflowr*^*20*^ website (https://phipsonlab.github.io/propeller-paper-analysis/), with the original source code available on github (https://github.com/phipsonlab/propeller-paper-analysis/tree/master). The *propeller* methods are available in the `speckle` R package (https://github.com/phipsonlab/speckle).

## 6. Materials and Methods

### 6.1 The *propeller* statistical model

Let *x*_*ij*_ denote the number of cells observed for cell type *i* for sample *j*, and let *N*_*j*_ denote the total number of cells observed for sample *j*. For each cell type *i* and sample *j*, we calculate the cell type proportions as

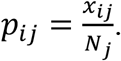

We perform a variance stabilising step in *propeller* by performing a data transformation. The arcsin square root transform, denoted *z*_*ij*_, is simply calculated as follows:

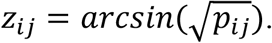

For the logit transform, more care is needed to deal with zeroes in the data. We thus add a count of 0.5 to *x*_*ij*_ when calculating the proportions:

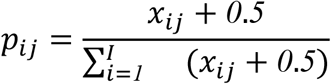

The logit transform is then given by

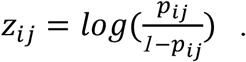

Let 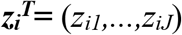 denote the vector of transformed proportions for cell type *i* for samples 1,…,*J*. We fit a linear model,

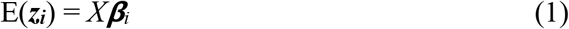

where *X* is a design matrix of full column rank and ***β***_*i*_ is a vector of coefficients. In general, ***β***_*i*_ can be estimated by

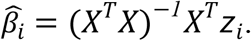

The variance of the transformed proportions for the *i*^*th*^ cell type is denoted *s*_*i*_^*2*^ and are the residuals obtained from fitting the linear model in (1). Moderated t-statistics for each contrast *k* are calculated as

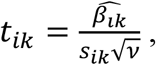

where *v* is the appropriate diagonal element from the positive definite matrix (*X*^*T*^*X*)^-1^, and *s*_*ik*_^*2*^ are the squeezed variances from Smyth (2004). The *t*_*ik*_ follow a scaled t distribution with augmented degrees of freedom *d*_*o*_*+d*_*i*_. Once p-values are obtained from the moderated t-statistics, they are adjusted for multiple testing using the method of Benjamini and Hochberg^15^, which has been shown to be robust to dependence of the test statistics^21^. The BH procedure is also robust to the number of cell types and will compute when the number of cell types is as low as two.

### 6.2 Estimating the parameters of the Beta distribution from observed cell type counts

We used the method of moments to estimate the parameters *α* and ***β*** of the Beta distribution from observed cell type counts *x*_*ij*_ from real single cell data in some of our simulated datasets. We implemented the estimation procedure on both cell type counts and proportions in the *estimateBetaParamsFromCounts* and *estimateBetaParam* functions respectively. We found the estimation procedure more stable on the cell type counts and it is described here.

The cell type proportions *p*_*ij*_ for cell type *i* and sample *j* are calculated as

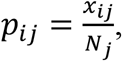

where

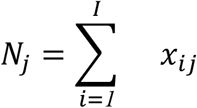

are the total cell counts for sample *j*. In practice the cell type counts are scaled to the median *N*_*j*_, such that all *N*_*j*_ =*N*. For cell type *i*, the mean and variance of the Beta distribution are given by

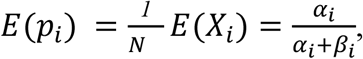

and

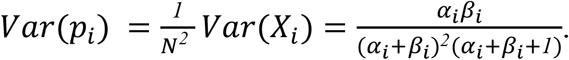

The *k*^*th*^ raw moment can be estimated from the *k*^*th*^ raw sample moment for cell type *i*

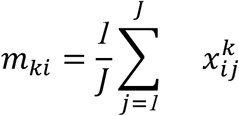

The estimated first and second moments for cell type *i* are given by

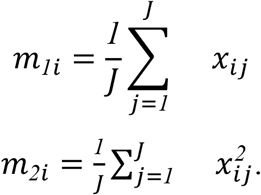

It can hence be shown through the mathematical relationships between the mean, variance and moments that estimators for *α*_*i*_ and ***β***_*i*_ can be obtained by

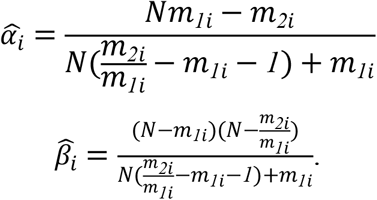

### 6.3 The hierarchical model for the simulated datasets

Let ***π*** denote a vector of true probabilities for the cell types. For a given value of ***α, β*** can be calculated as

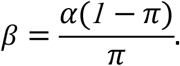

Alternatively, ***α*** and ***β*** can be estimated from observed cell type counts in real single cell datasets using the method of moments described above.

The sample proportions *p*_*ij*_ for cell type *i* and sample *j* are generated from a Beta distribution:

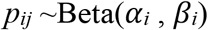

The cell type counts *x*_*ij*_ are sampled from a binomial distribution

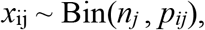

where the total number of cells per sample *j* are sampled from a negative binomial distribution

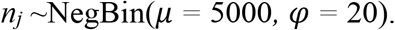

The resulting cell type counts follow a beta-binomial distribution.

### 6.4 Implementation of the different methods in R

For the chi-square test, cells from the samples within each group were summed and the *prop*.*test* function was used to obtain p-values for each cell type. For the negative binomial models, the `edgeR` Bioconductor package was used. For the classic negative binomial regression, the *glmFit* and *glmLRT* functions were used to model cell type counts. For quasi-likelihood negative binomial, *glmQLFit* and *glmQLFTest* were used. To fit Poisson regression, the *glmFit* and *glmLRT* functions in `edgeR` were used with the dispersion set to zero. For beta-binomial, an alternate parameterisation for the negative binomial was employed, as described in Chen et al.,^22^ to analyse bisulfite sequencing data. For logistic binomial regression, the same approach was taken but with the dispersion set to zero. We implemented the CODA method from first principles. First, we replaced any zeroes in the data with 0.5. For each sample *j* we calculated the geometric mean. The centred log-ratio transformation divides the observed cell type count for sample *j* by the geometric mean for sample *j* and then takes the log of the ratio. Normal regression theory can then be applied to the transformed data. We used `limma` to fit linear regression models on the transformed data.

## Acknowledgements

We thank Xinyi Jin for preparing the `speckle` package for submission to Bioconductor. We thank Dr Jovana Maksimovic for suggesting the COVID-19 dataset as an application dataset, and Prof Gordon Smyth for discussions regarding multiple testing adjustment under dependence of test statistics.

## References

1. Zeisel, A. et al. Brain structure. Cell types in the mouse cortex and hippocampus revealed by single-cell RNA-seq. Science 347, 1138–1142 (2015).

2. Bornstein, C. et al. Single-cell mapping of the thymic stroma identifies IL-25-producing tuft epithelial cells. Nature 559, 622–626 (2018).

3. Liu, Y. et al. Single-cell RNA-seq reveals the diversity of trophoblast subtypes and patterns of differentiation in the human placenta. Cell Res. 28, 819–832 (2018).

4. Combes, A. N. et al. Single cell analysis of the developing mouse kidney provides deeper insight into marker gene expression and ligand-receptor crosstalk. Development 146, (2019).

5. Sim, C. B. et al. Sex-Specific Control of Human Heart Maturation by the Progesterone Receptor. Circulation 143, 1614–1628 (2021).

6. Huang, Z. et al. Effects of sex and aging on the immune cell landscape as assessed by single-cell transcriptomic analysis. Proc. Natl. Acad. Sci. U. S. A. 118, (2021).

7. Ren, X. et al. COVID-19 immune features revealed by a large-scale single-cell transcriptome atlas. Cell 184, 1895–1913.e19 (2021).

8. Bunis, D. G. et al. Single-Cell Mapping of Progressive Fetal-to-Adult Transition in Human Naive T Cells. Cell Rep. 34, 108573 (2021).

9. Xu, J. et al. Genotype-free demultiplexing of pooled single-cell RNA-seq. Genome Biol. 20, 290 (2019).

10. Huang, Y., McCarthy, D. J. & Stegle, O. Vireo: Bayesian demultiplexing of pooled single-cell RNA-seq data without genotype reference. Genome Biol. 20, 273 (2019).

11. Crowell, H. L. et al. muscat detects subpopulation-specific state transitions from multi-sample multi-condition single-cell transcriptomics data. Nat. Commun. 11, 6077 (2020).

12. Tan, Q. et al. Handling blood cell composition in epigenetic studies on ageing. International journal of epidemiology vol. 46 1717–1718 (2017).

13. Smyth, G. K. Linear models and empirical bayes methods for assessing differential expression in microarray experiments. Stat. Appl. Genet. Mol. Biol. 3, Article3 (2004).

14. Efron, B. & Morris, C. Stein’s Paradox in Statistics. Sci. Am. 236, 119–127 (1977).

15. Benjamini, Y. & Hochberg, Y. Controlling the False Discovery Rate: A Practical and Powerful Approach to Multiple Testing. J. R. Stat. Soc. Series B Stat. Methodol. 57, 289–300 (1995).

16. Orchestrating Single-Cell Analysis with Bioconductor. https://bioconductor.org/books/release/OSCA/.

17. Quinn, T. P. et al. A field guide for the compositional analysis of any-omics data. Gigascience 8, (2019).

18. Liao, M. et al. Single-cell landscape of bronchoalveolar immune cells in patients with COVID-19. Nat. Med. 26, 842–844 (2020).

19. Simmons, S. Cell Type Composition Analysis: Comparison of statistical methods. bioRxiv 2022.02.04.479123 (2022) doi:10.1101/2022.02.04.479123.

20. Blischak, J. D., Carbonetto, P. & Stephens, M. Creating and sharing reproducible research code the workflowr way. F1000Res. 8, 1749 (2019).

21. Benjamini, Y. & Yekutieli, D. The Control of the False Discovery Rate in Multiple Testing under Dependency. Ann. Stat. 29, 1165–1188 (2001).

22. Chen, Y., Pal, B., Visvader, J. E. & Smyth, G. K. Differential methylation analysis of reduced representation bisulfite sequencing experiments using edgeR. F1000Res. 6, 2055 (2017).

