## Supplementary Material for "*propeller*: testing for differences in cell type proportions in single cell data"

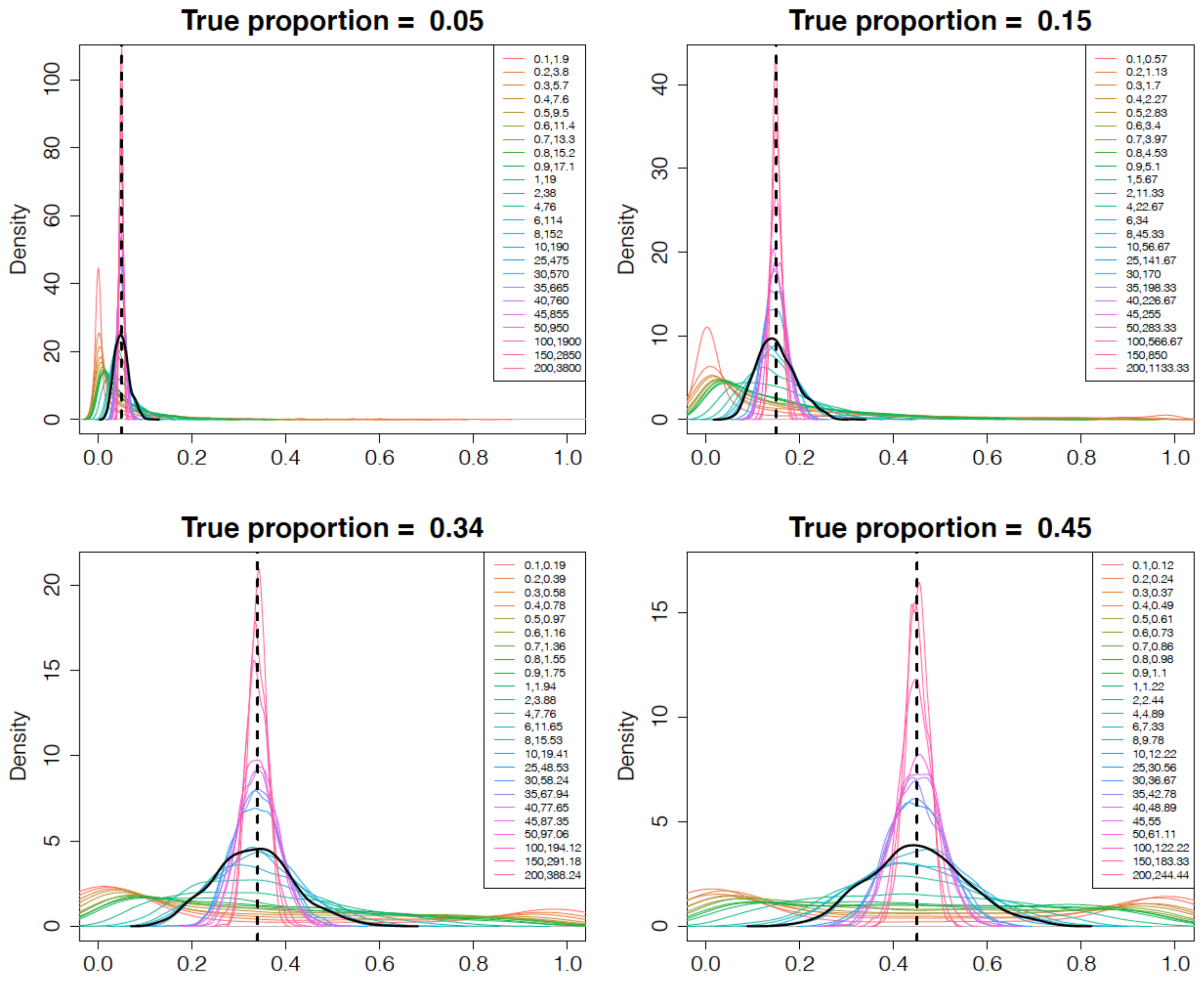


**Supplementary Figure 1. Hyperparameters for Beta distribution in the simulation setting.** The values of alpha and beta determine how diffuse the prior is under the Beta Binomial hierarchical model assumed in the simulation. The two values in the legend correspond to a range of values for alpha and beta. The density plots are based on 1000 sampled data points from a Beta distribution with corresponding alpha and beta parameter values. The line in black highlights the case where alpha = 10.

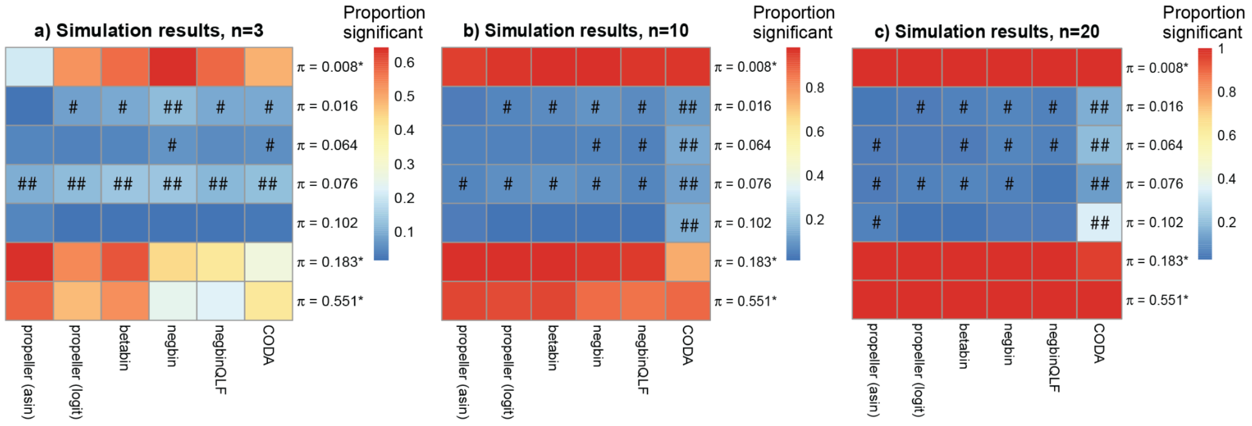


**Supplementary Figure 2. Power to detect true differences in abundance in 1000 simulated datasets.** We simulated cell type differences between two groups for three cell types, with the remaining four cell types not changing between the two groups. We varied the sample size between two groups: $n = 3$, 10, and 20. For each of the seven cell types, we calculated the proportion of simulated datasets with p-value < 0.05 for each of the six models. As sample sizes increase, all methods detect the true positive cell types in a higher proportion of simulated datasets. For the true negatives, dark blue without the # symbol indicates good false discovery rate with proportion significant <0.05, # indicates proportion significant between 0.05 and 0.1, and ## indicates proportion significant > 0.1. As the sample size increases, the number of false discoveries for the CODA method dramatically increases.

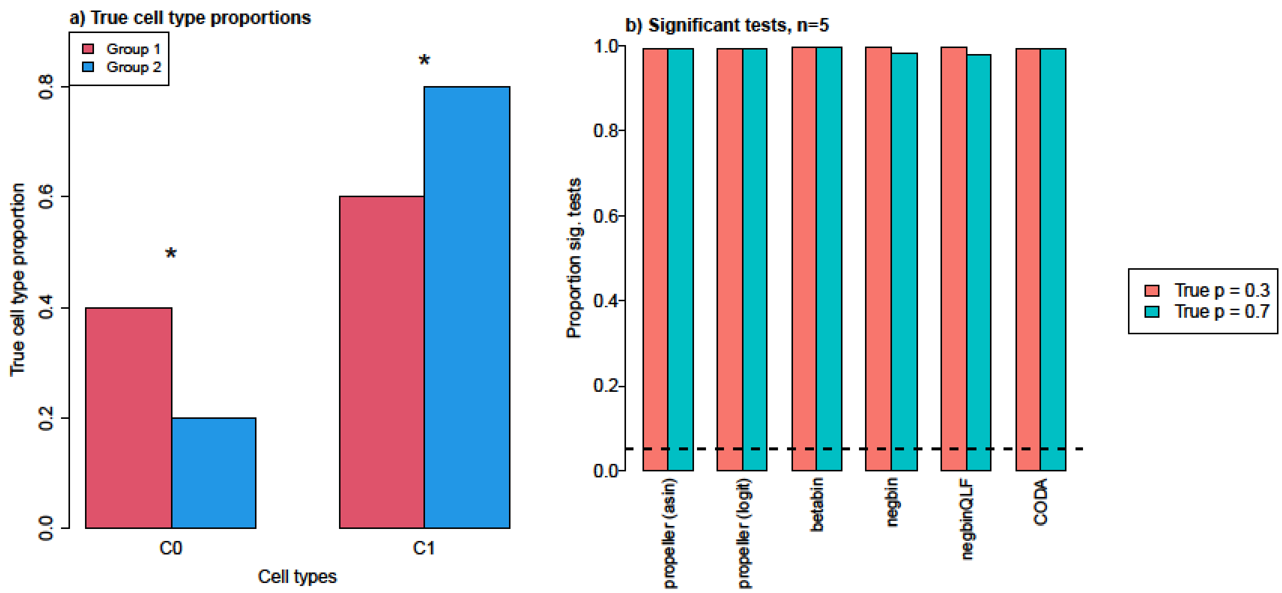


**Supplementary Figure 3. Special case: two cell types and true differences between two groups.**

**a.** True cell type proportions for the simulation model.

**b.** Proportion of significant tests for each cell type in the 1000 simulated datasets for the six different methods. In the majority of datasets, both cell types were detected as significantly different (p-value < 0.05) between Group 1 and Group 2 by all methods. There was a slight decrease in power for the negative binomial methods for the more abundant cell type (cell type C1).

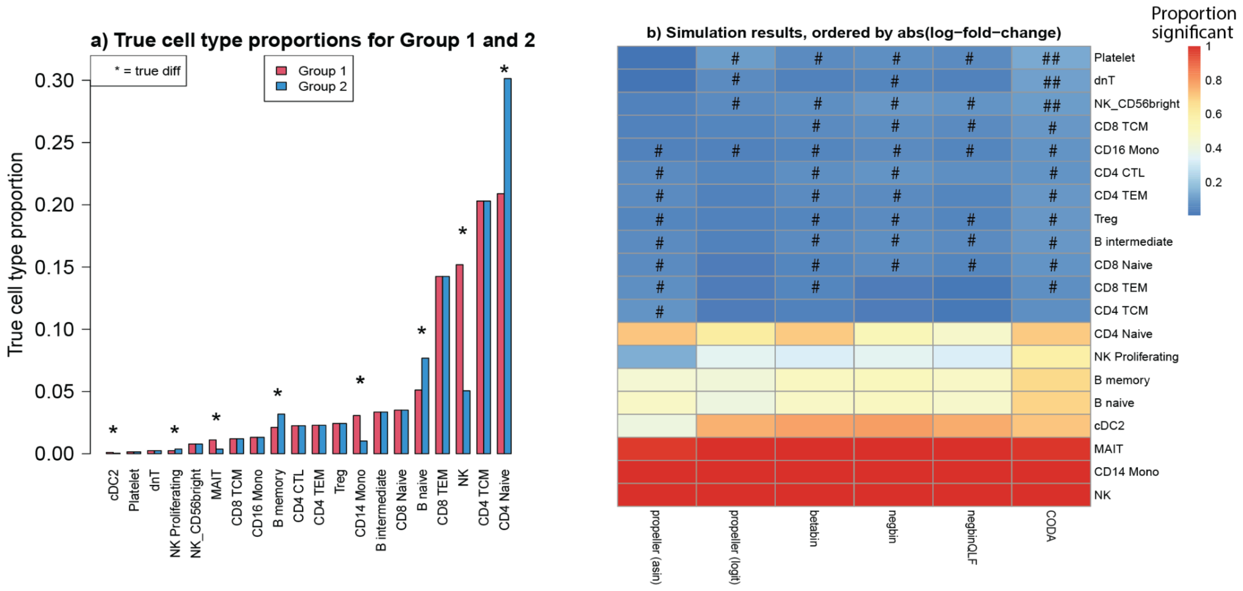


**Supplementary Figure 4. Special case: 20 cell types and true differences between two groups.**

**a.** True cell type proportions for group 1 and group 2 for the simulation model. The 8 cell types denoted with * have differences in abundance between group 1 and group 2 that range between 1.4 and 3 fold. The remaining 12 cell types do not change between groups.

**b.** Heatmap showing proportion of simulated datasets with p-value < 0.05 for each of the six models for the 20 different cell types. The cell types are ordered by the absolute value of the log-fold changes such that the top 12 cell types are not changing, and the bottom 8 change, with NK cells having the biggest change in abundance between Group 1 and Group 2. We see that generally the methods all perform well when the log-fold changes are larger. For smaller log-fold changes, there is reduced power to detect significant changes in abundance. The CODA method generally has good power to detect the true positives, but has more false discoveries than the other methods. propeller(logit) outperforms propeller(asin) and has the best false discovery rate control, and the negative binomial methods perform comparably to the other methods in this scenario. For the true negatives, dark blue without the # symbol indicates good false discovery rate with proportion significant <0.05, # indicates proportion significant between 0.05 and 0.1, and ## indicates proportion significant > 0.1. The results shown are based on 1000 simulated datasets.

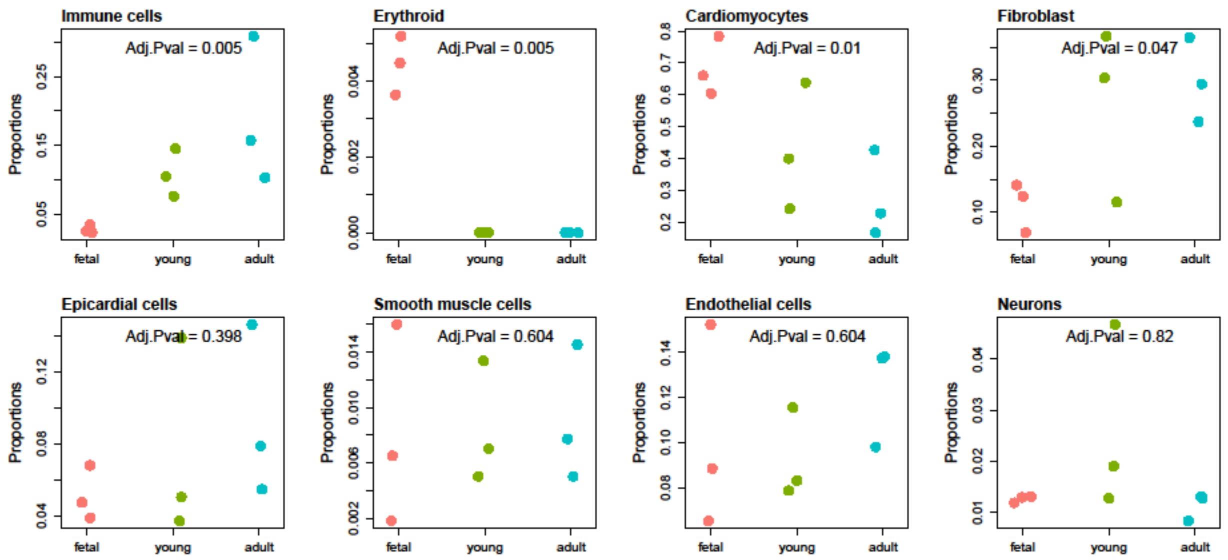


**Supplementary Figure 5.** Cell type proportions for each cell type in human heart single nuclei RNA-seq dataset. The cell types are ordered from most significant (Immune cells) to least significant (Neurons). The first four cell types are statistically significant at FDR < 0.05.

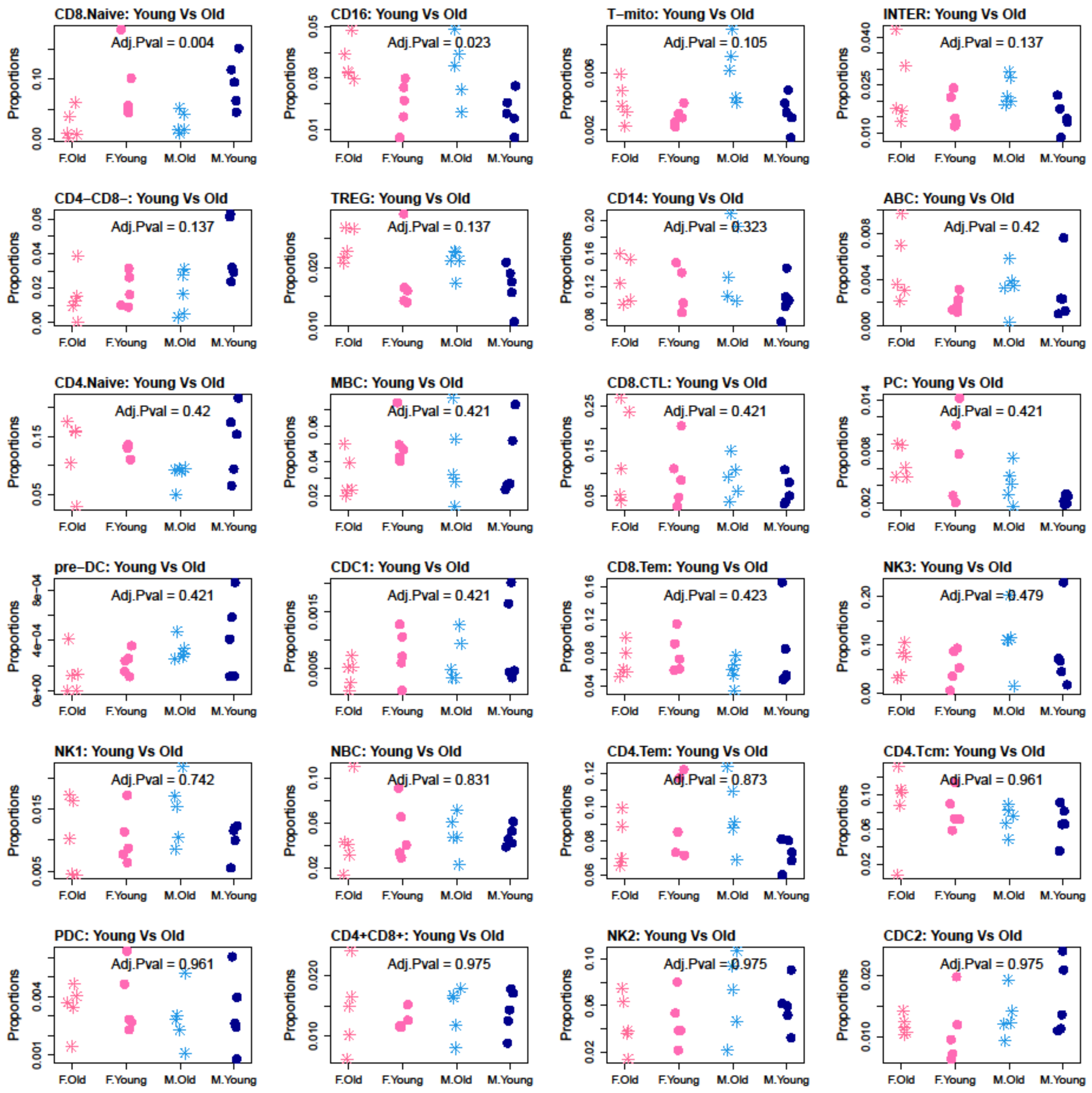


**Supplementary Figure 6.** Cell type proportions for each cell type for the PBMC ageing and sex single cell data. The cell types are ordered by statistical significance when testing Young vs Old, taking sex into account. The first two cell types, CD8 naive and CD16, are statistically significant at FDR < 0.05, the remaining cell types do not achieve statistical significance.

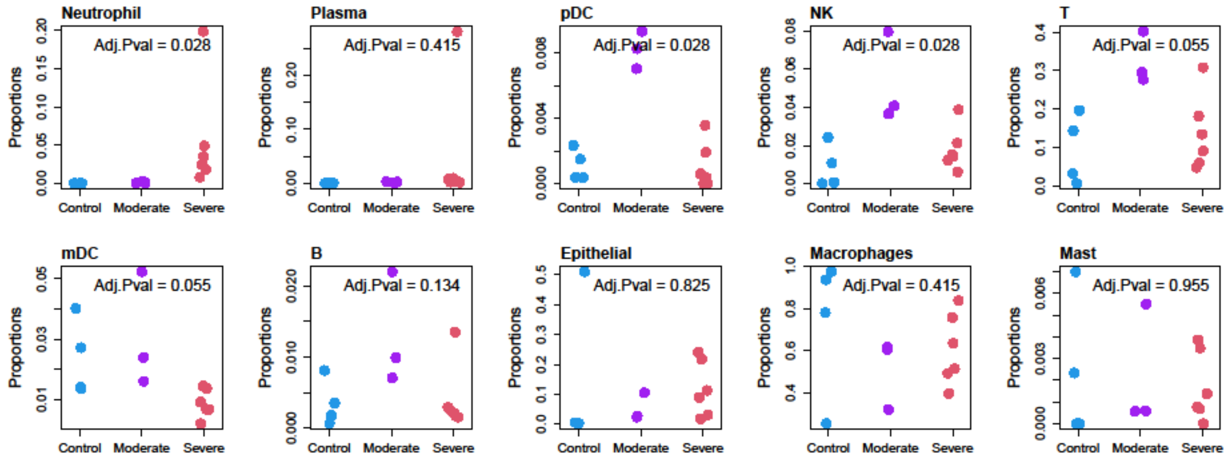


**Supplementary Figure 7.** Cell type proportions for each cell type in COVID-19 scRNA-seq dataset. The cell types are ordered from most significant (Neutrophil) to least significant (Mast) when analysed using propeller(logit). The first four cell types are statistically significant at FDR < 0.05 using propeller(logit). The adjusted p-values printed on the plots are from propeller(asin). Plasma appears to have an outlier sample in the severe COVID-19 group, and when the analysis is re-run with propeller(asin), this cell type is no longer significant.
